# A TurboID-based proximity labelling method for detecting extracellular protein interactions in plants

**DOI:** 10.64898/2026.09.22.753433

**Authors:** Abdelrahman M Qutb, Camille-Madeleine Szymansky, Graeme J Kettles

## Abstract

The apoplast is a primary location for interactions between plants and invasive pathogens and is the site where many pathogen-secreted effectors are perceived by cell surface immune receptors to activate defence responses. However, our understanding of apoplastic interactions remains limited because protein interactions are difficult to investigate in this harsh extracellular environment. TurboID-based proximity labelling (PL) has emerged as a powerful approach for studying protein interactions in plants, though its application has to date been restricted to intracellular proteins. Here, we designed and validated a TurboID-based PL strategy for investigating protein interactions in the leaf apoplast using the well-characterised interaction between the *Phytophthora infestans* elicitor INF1 and the receptor-like protein (RLP) REL in *Nicotiana benthamiana*. Transient expression of SP-INF1-TurboID (INF1-T) induced a cell death (CD) response comparable to that triggered by native INF1, demonstrating that fusion of the TurboID tag did not impair INF1 recognition by REL. Apoplastic localisation of both INF1-T and the control construct SP-eGFP-TurboID (eGFP-T) was confirmed, validating their suitability for PL experiments. Efficient TurboID-mediated biotinylation was achieved in the apoplast using co-infiltration of biotin, ATP and magnesium acetate. Streptavidin-HRP immunoblotting revealed distinct biotinylation profiles for INF1-T and eGFP-T. Furthermore, co-immunoprecipitation demonstrated specific biotinylation of REL by INF1-T, but not by eGFP-T, in wild-type, *bak1* and *sobir1*/*sobir1*-like *N. benthamiana*. These findings demonstrate that TurboID-based PL is functional in the apoplast and provides a proof-of-concept for investigating elicitor-receptor interactions in this compartment.

## Introduction

Plant responses to biotic stresses are mediated through a range of pathways, where interactions between different classes of molecules from both partners play critical roles (Struk et al. 2019). Among these, protein–protein interactions (PPIs) between host and pathogen proteins are a core determinant on the outcome of the interaction (Lampugnani et al. 2018). Several methods have been developed to investigate PPIs, among which *in planta* expression systems utilise the cellular environment to study these interactions (Cuadrado and Van Damme 2024). These approaches rely on plant-based expression and offer the advantage of examining PPIs within their native context. Co-immunoprecipitation (Co-IP) is a widely used approach for investigating PPIs. In this technique, a bait protein is captured from a total protein extract using an immobilised antibody, and potential interacting partners are subsequently identified through Western blotting or mass spectrometry (MS; Masters 2004). Despite its value, the method has limitations, as it primarily detects strong and stable direct interactions (Struk et al. 2019), making it less suitable for transient, weak, or proximal interactions. Another biochemical strategy for PPIs is affinity purification combined with MS (AP-MS; Struk et al., 2019). This approach also has drawbacks, as it is not well suited to detecting proteins that interact only weakly or transiently with the target protein (Zhang et al. 2019). A more recent approach uses enzyme-catalysed labelling termed proximity labelling (PL; Kim & Roux, 2016). In this method, the bait protein is fused to a promiscuous labelling enzyme that marks nearby proteins once a specific substrate is supplied (Kim & Roux, 2016). The labelled candidates can then be isolated through affinity purification and subsequently characterised by MS. For PL, two enzymes have been employed: soybean ascorbate peroxidase (APEX2) and a promiscuous variant of the *Escherichia coli* biotin ligase (BioID; Cuadrado and Van Damme, 2024). Each, however, has potential drawbacks. APEX2 depends on hydrogen peroxide (H_2_O_2_) to initiate labelling, which poses toxicity issues for living material (Lam et al. 2014). BioID avoids this problem, but it’s slow catalytic rate makes it less effective for detecting transient interactions (Roux et al. 2012). To address these limitations, an improved enzyme, TurboID, was developed. Here, an evolved *E. coli* biotin ligase BirA was created by yeast display to create highly promiscuous labelling variants. TurboID enables rapid, non-toxic PL within just 10 mins. The result is a fast and non-toxic enzyme suitable for use across diverse organisms and for studying dynamic cellular processes (Branon et al. 2018). This biotin ligase functions without H_2_O_2_ and is approximately one hundred times faster than BioID (Branon et al. 2018). At present, TurboID represents the most widely adopted PL system and has been successfully applied in diverse plant models to investigate PPIs (Zhang et al. 2022). TurboID-based PL has been successfully applied to investigate intracellular interactions, such as signalling networks involving the brassinosteroid insensitive (BIN2) kinase and the regulation of nucleotide-binding leucine-rich repeat receptors (NLRs; Zhang et al. 2019). However, to our knowledge, it has not yet been effectively utilised to study extracellular immune interactions (Kim et al. 2023). Apoplastic interactions play a role in all pathosystems but are particularly relevant for pathogens that remain extracellular throughout their life cycle. For example, the wheat pathogen *Zymoseptoria tritici* secretes apoplastic effectors that modify host processes such as cell death, ROS production and immune signalling, but are subject to recognition by host cell surface receptor-like kinases (RLKs) (Welch et al. 2022; Qutb et al. 2024; Zhong et al. 2017; Saintenac et al. 2021).

In this study, we developed a TurboID-based PL approach for studying receptor– ligand interactions in the leaf apoplast. As the use of TurboID for studying protein interactions in the apoplast has not yet been established, we validated the approach using known interactions between pathogen elicitors and their cognate cell surface receptors. Specifically, we used the interactions between *Phytophthora sojae* XEG1 and the cell-surface receptor-like protein (RLP) RXEG1 (Wang et al. 2018), and the *P. infestans* elicitor INF1 and the cell-surface RLP REL (Chen et al. 2023), using the *N. benthamiana* transient expression system.

## Results

### TurboID tagging does not compromise INF1-induced cell death (CD)

Following cloning of the PL constructs (Figure 1A), recombinant proteins were Agroexpressed in *N. benthamiana* leaves and total protein extracted at 48 hours post infiltration (hpi) with constructs expressed from either pEAQ-HT-DEST3 (pEAQ3; Sainsbury et al., 2009) or and pEARLYGATE101 (pEG101; Earley et al., 2006). Western blots using an α-Myc–HRP antibody showed that protein accumulation was consistently higher from pEAQ3 than from pEG101 (Figure 1B). In particular, SP-eGFP-TurboID (eGFP-T) accumulated to substantially higher levels when expressed from pEAQ3 than from pEG101. Furthermore, SP-XEG1-TurboID (XEG1-T) and SP-INF1-TurboID (INF1-T) were detected only when expressed from pEAQ3, whereas eGFP-T accumulated to markedly higher levels than either XEG1-T or INF1-T. In contrast, SP-INF1 (INF1) was not detected when expressed from either vector (Figure 1B).

**Figure 1.**
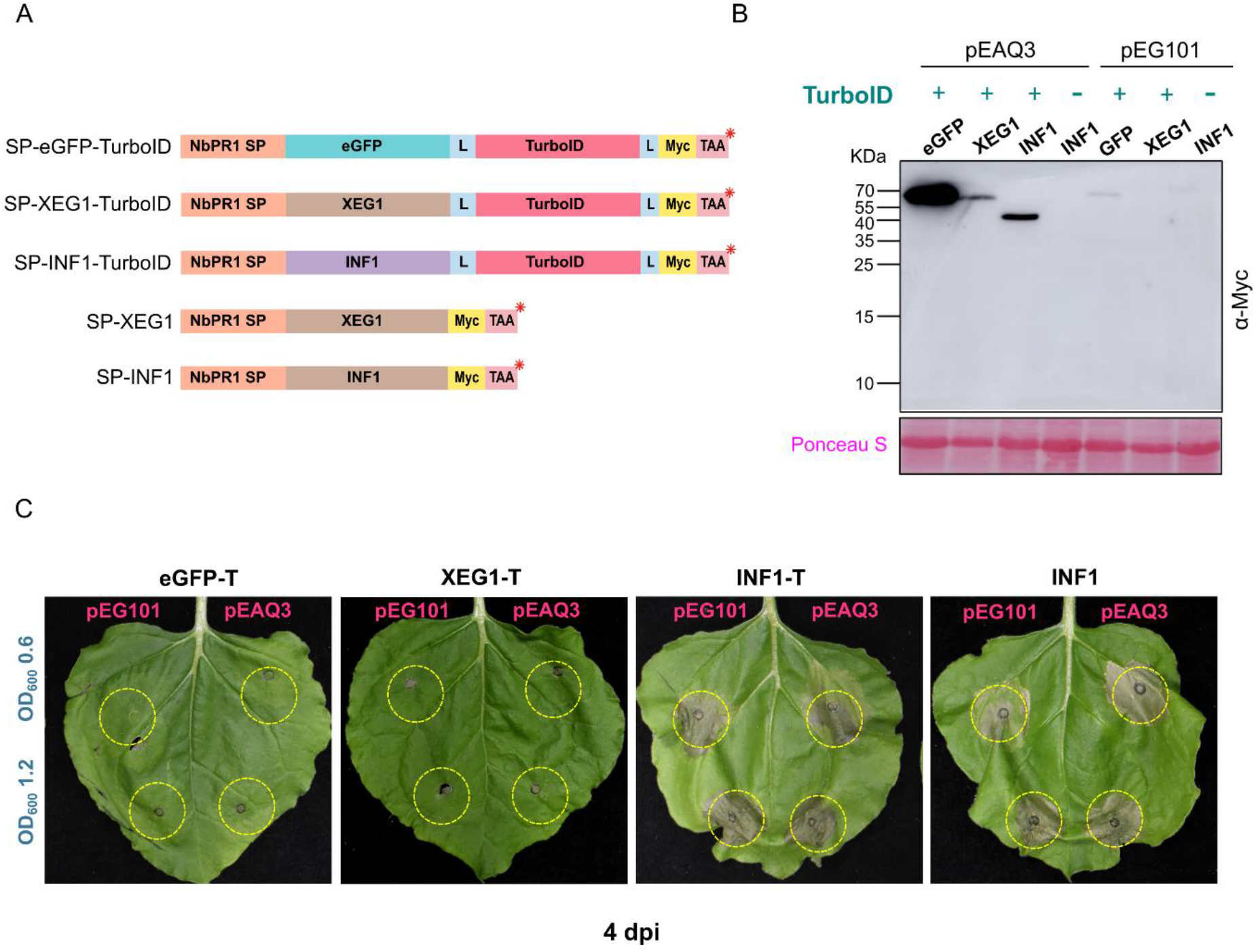
CD response of PL constructs in *N. benthamiana* WT leaves. (A) Schematic of the PL constructs used in this study. (B) Accumulation of the PL constructs expressed from the pEAQ3 and pEG101 vectors in *N. benthamiana*. Protein accumulation was analysed by western blot using α-Myc-HRP. Each lane represents a different PL construct expressed from the indicated vector. Equal protein loading indicated by Ponceau S staining. (C) Plant phenotypes induced by *Agrobacterium*-mediated expression of PL constructs. Cell density at OD_600_ indicated. Leaves were assessed at 4 dpi. The experiment was repeated twice with similar results.

In parallel, the CD-inducing activity of the PL constructs was assessed. At 4 days post-infiltration (dpi), both INF1 and INF1-T induced obvious CD when expressed from either pEAQ3 or pEG101 (Figure 1C). In contrast, XEG1-T did not induce CD when expressed from either vector. As expected, eGFP-T did not induce CD, indicating that apoplast-targeted TurboID alone does not trigger a CD response (Figure 1C). In a further experiment, transient expression of SP-XEG1 (XEG1) lacking TurboID in WT plants also failed to induce CD (Figure S1). These results demonstrate that fusion of TurboID to INF1 did not compromise its ability to induce CD, confirming that the TurboID-tagged construct retained its biological activity *in planta*.

### Optimisation of eGFP-T and INF1-T protein accumulation

Due to absence of CD and low accumulation of XEG1 PL constructs, subsequent experiments focused on eGFP and INF1 interactions. As INF1 constructs accumulated to low levels and triggered CD in WT plants, it was necessary to achieve comparable protein accumulation between eGFP-T and INF1-T in WT plants. This would aid identification of specific labelling in future biotinylation assays. Therefore, the accumulation of both constructs was investigated by testing different infiltration densities (OD_600_) of the *Agrobacterium* strains across four time points (Figure 2). In this experiment, three OD_600_ values were assessed for both eGFP-T and INF1-T (0.1, 0.4, and 0.8). At 24 hpi, neither eGFP-T nor INF1-T was detectable at any of the three OD_600_ values (Figure 2). Importantly, the abundance of both constructs increased in line with higher OD_600_ densities of the *Agrobacterium* strains (Figure 2). eGFP-T abundance continued to rise gradually over the later time points, whereas INF1-T abundance increased only up to 60 hpi. The reduction at 72 hpi time point was likely due to CD development in agroinfiltrated leaves, which negatively affected protein accumulation (Figure 2). Based on these results, eGFP-T expressed at OD_600_ 0.4 was found to reach a similar abundance to INF1-T expressed at OD_600_ 0.8 at 48 hpi (Figure 2).

**Figure 2.**
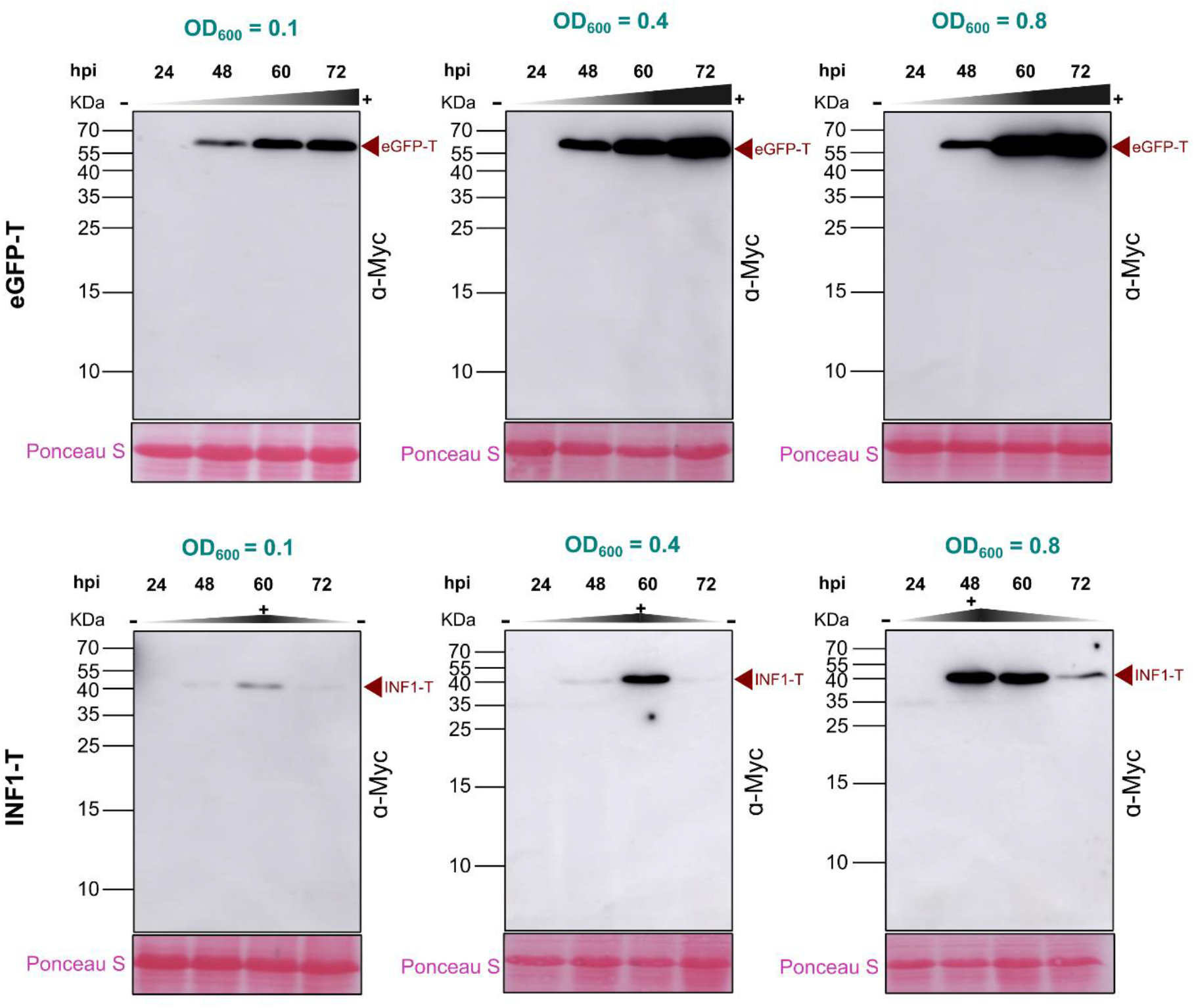
Optimisation of eGFP-T and INF1-T protein accumulation in *N. benthamiana*. Protein accumulation of eGFP-T and INF1-T expressed from pEAQ3 assessed at three *Agrobacterium* OD_600_ densities (0.1, 0.4, 0.8) and four time points (24, 48, 60 and 72 hpi) following *Agrobacterium*-mediated expression. Protein accumulation detected by Western blot using α-Myc-HRP. Equal protein loading indicated by Ponceau S staining.

The CD response triggered by INF1-T reduced protein accumulation at later time points, limiting downstream biochemical analyses. Therefore, several approaches were investigated to weaken or abolish this response. To determine whether INF1-T-triggered CD was light-dependent, agroinfiltrated plants were maintained in the dark for 3 dpi. However, CD development was comparable between plants maintained in the dark and those grown under normal light conditions (Figure S2), indicating that INF1-T recognition and CD development are not light-dependent.

As calcium is a key regulator of plant CD signalling (Orrenius et al. 2015), the calcium channel blocker lanthanum chloride (LaCl_3_) was evaluated as a CD inhibitor. A 1 mM LaCl_3_ solution was co-infiltrated with *Agrobacterium* carrying the INF1-T construct. At 4 dpi, LaCl_3_ treatment did not reduce the CD phenotype compared with the untreated control (Figure S3A). Moreover, the LaCl_3_ treatment reduced INF1-T protein accumulation relative to the untreated control (Figure S3B), making this approach unsuitable for subsequent experiments.

As neither darkness nor LaCl_3_ treatment suppressed the INF1-T-induced CD response, a genetic approach was investigated. Since recognition of INF1 by the RLP REL requires the co-receptor kinases NbBAK1 and NbSOBIR1 (Chaparro-Garcia et al. 2011; Domazakis et al. 2018), *bak1* (Sun et al. 2022) and *sobir1* & *sobir1-like* (Huang et al. 2021) knockout mutants were tested for development of CD. Agroexpression of INF1-T in both mutant backgrounds abolished the CD phenotype observed in WT plants (Figure S4). Western blots showed that INF1-T accumulated progressively in both *bak1* and *sobir1/sobir1-like* plants up to 72 hpi (Figure S5), exhibiting a pattern similar to that of eGFP-T expression in WT plants (Figure 2). This indicated that removal of co-receptors can enhance TurboID-tagged elicitor accumulation through the absence of CD.

### eGFP-T and INF1-T localise to the apoplast and mediate protein biotinylation

As all PL constructs contained the *N. benthamiana* PR1a signal peptide (NbPR1 SP), their localisation to the apoplast was investigated after confirming their expression *in planta*. Confocal microscopy showed that eGFP-T fluorescence was confined to the cell periphery, consistent with apoplastic localisation, whereas the cytosolic GFP control was distributed throughout the cytoplasm (Figure S6A).

To confirm apoplastic localisation, INF1-T was expressed in *sobir1/sobir1-like* plants to prevent CD and facilitate protein accumulation up to 72 hpi. Apoplast wash fluid (AWF) was collected from leaves expressing eGFP-T or INF1-T and concentrated by spin column. Western blotting detected eGFP-T in the unconcentrated AWF, whereas INF1-T was not detected in either concentrated or unconcentrated samples (Figure S6B). However, re-infiltration of both concentrated and unconcentrated INF1-T AWF into WT plants induced CD (Figure S6C), indicating that active INF1-T was present in the AWF despite either (i) being below the detection limit of western blotting or (ii) cleavage of the Myc tag.

Having confirmed the apoplastic localisation of eGFP-T and demonstrated the presence of INF1-T in the apoplast, the ability of TurboID to biotinylate apoplastic proteins was investigated. Following eGFP-T and INF1-T expression, total protein was extracted at 48 hpi, following a second infiltration at 45 hpi which supplied biotin, ATP and Mg^2+^ to the apoplastic space. This allowed a 3 h window for the biotinylation reaction. Owing to the lower accumulation of INF1-T, five biological replicates were analysed by western blot and the three samples with the highest protein accumulation (samples 1, 2 and 5) were selected for subsequent analyses (Figure S7). Both eGFP-T and INF1-T samples produced strong biotinylation signals, demonstrating that TurboID was active in the apoplast in the presence of supplied cofactors (Figure 3). Notably, the biotinylation profiles of eGFP-T and INF1-T were distinct, indicating that the two constructs labelled different sets of proteins. Probing with α-Myc–HRP confirmed enrichment of both eGFP-T and INF1-T, consistent with self-biotinylation of the TurboID fusion proteins (Figure 3).

**Figure 3.**
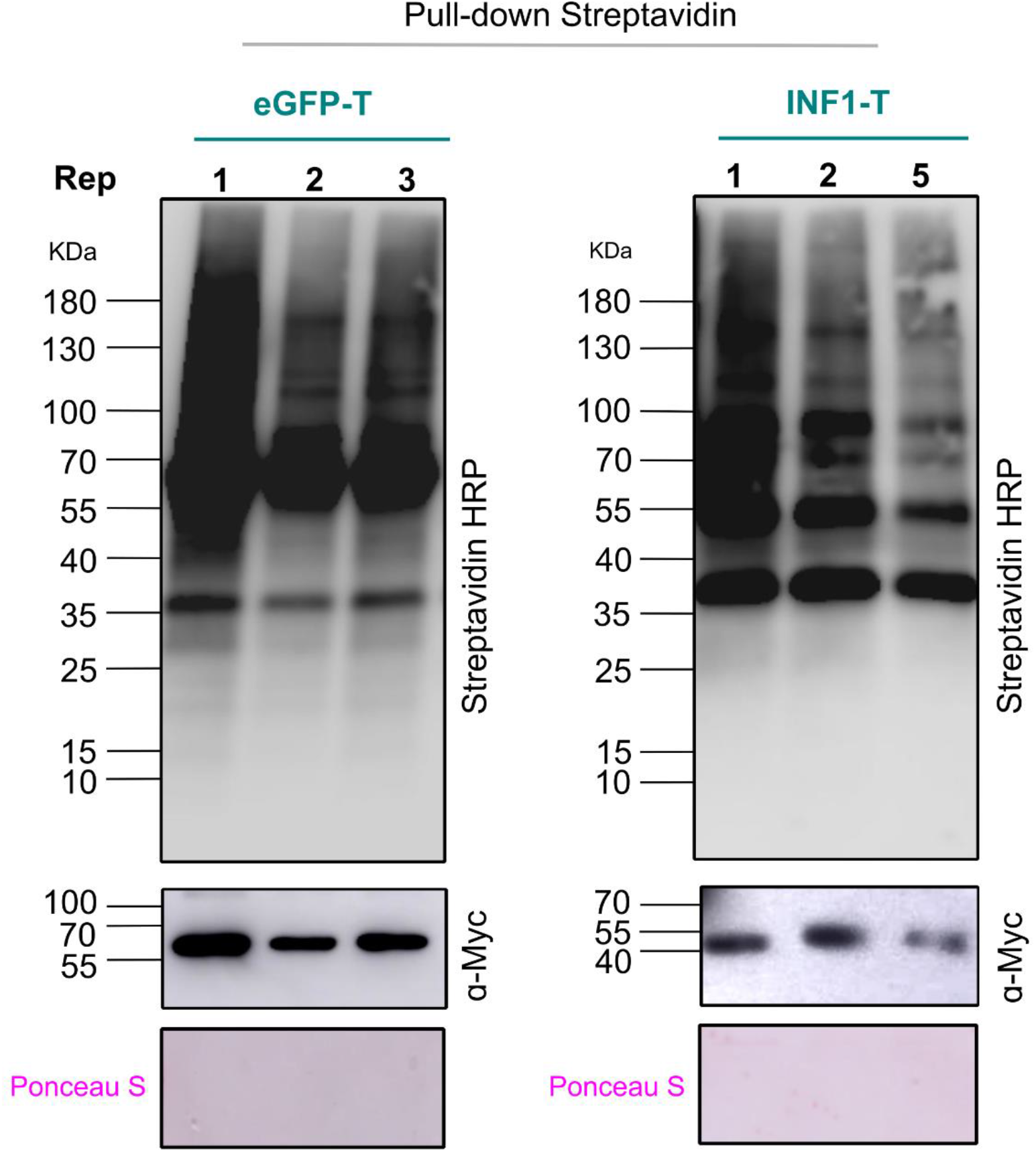
TurboID-based biotinylation in *N. benthamiana* apoplast. Protein biotinylation profiles generated by INF1-T and eGFP-T expression in the *N. benthamiana* apoplast. Three biological replicates shown per treatment. Lower panels indicate self-biotinylation of INF1-T and eGFP-T. Equal protein loading indicated by Ponceau S staining following protein enrichment by streptavidin bead purification.

### REL is specifically biotinylated by INF1-T in the apoplast

To assess whether INF1-T specifically labels the receptor REL, the CD response following co-expression of 35S-NbREL-GFP (REL-GFP) with the PL constructs was assessed and quantified in three genetic backgrounds. Expression of REL-GFP alone did not induce CD in WT plants (Figure 4A). In contrast, co-expression of REL-GFP with INF1 or INF1-T induced CD in WT plants (Figure 4A), whereas no CD was observed in *sobir1/sobir1-like* plants (Figure 4B). In *bak1* plants, co-expression of REL-GFP with INF1 resulted in a partial CD response. Quantification of CD in *bak1* plants showed a significant increase in CD induced by REL-GFP co-expressed with INF1-T compared with eGFP-T (Student’s *t*-test, *P* < 0.01; Figure 4C), whereas expression of the INF1 constructs alone did not induce CD.

**Figure 4.**
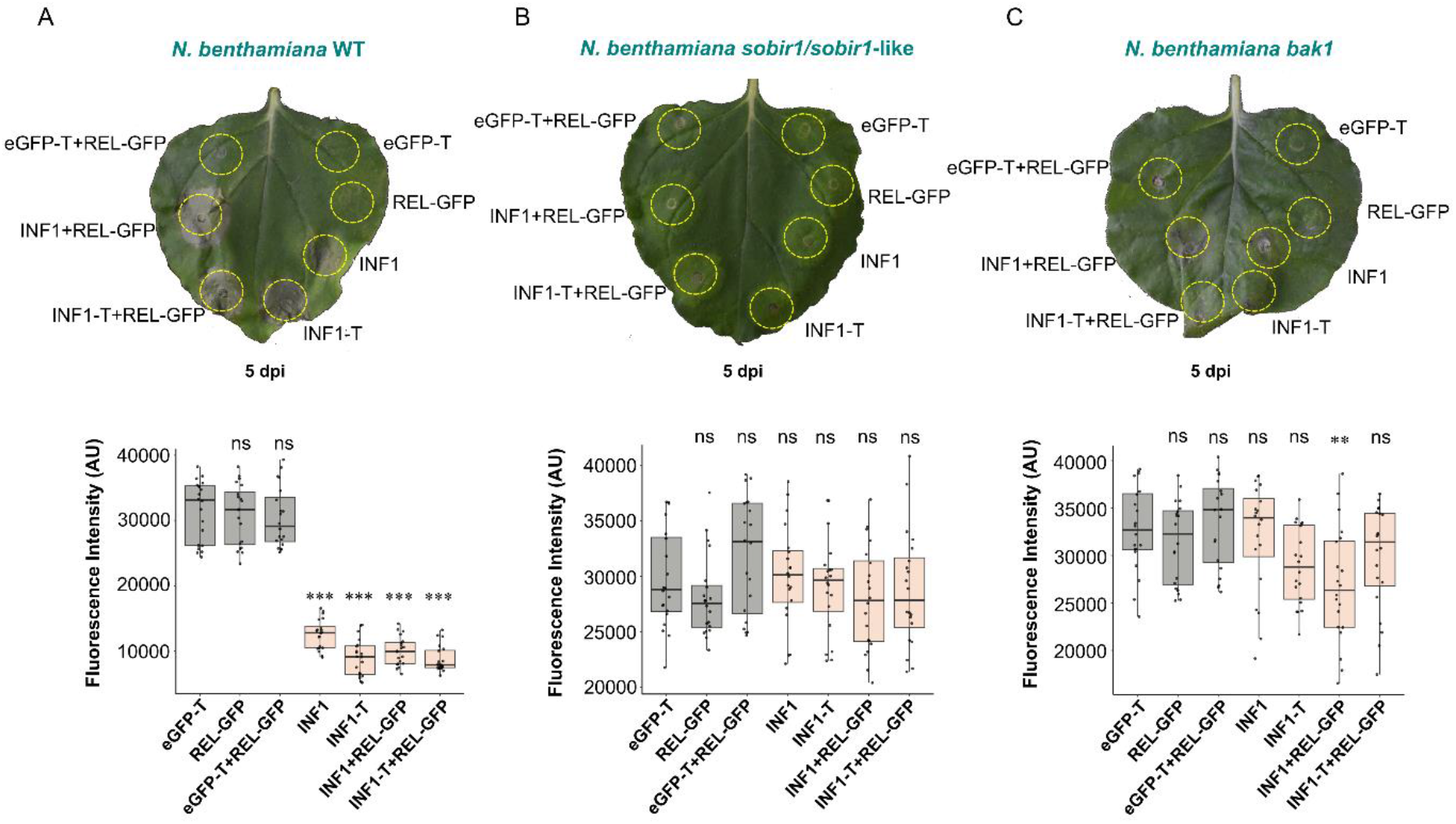
CD phenotypes of REL-GFP coexpression with PL constructs. (A) CD phenotypes induced in (A) *N. benthamiana* WT, (B) *sobir1/sobir1-like* and (C) *bak1*. CD was quantified by red fluorescence imaging (arbitrary units, AU) using 20 leaves per treatment. Light orange boxes indicate treatments expressing INF1 or INF1-T. Statistical significance was determined using Student’s *t*-test (\*\**P* < 0.01, \*\*\**P* < 0.001; ns, not significant). CD was assessed at 5 dpi.

To determine whether REL was specifically biotinylated by INF1-T, a Co-IP approach was employed. As REL overexpression accelerated CD development in WT plants, samples were collected at 36 hpi. To allow greater protein accumulation before the onset of CD, samples were collected at 46 hpi in *bak1* plants and at 60 hpi in *sobir1/sobir1-like* plants, in which CD was absent. Across the input samples, eGFP-T was detected in all plants (Figure 5A–C), whereas INF1-T and INF1 were not detected in WT, likely because of lower abundance and earlier sampling time (Figure 5A). However, INF1-T was detected in both *bak1* and *sobir1/sobir1-like* plants (Figure 5B, C). Immunoblotting with α-GFP–HRP detected eGFP-T in all three genetic backgrounds, whereas REL-GFP was not detected in the input samples (Figure 5A– C).

**Figure 5.**
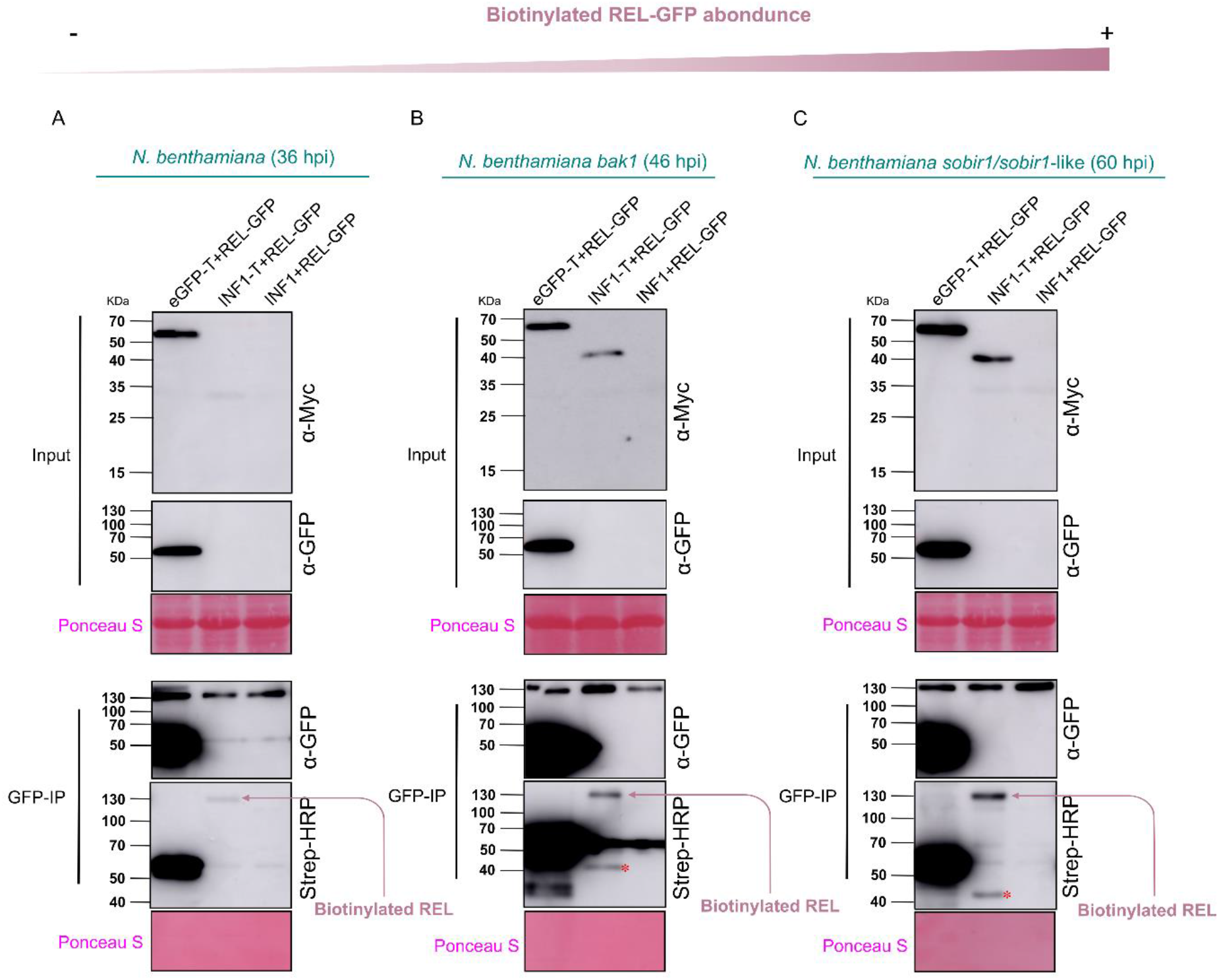
Apoplastic biotinylation of REL-GFP by INF1-T. Coexpression of REL-GFP and TurboID constructs in (A) WT, (B) *bak1* and (C) *sobir1/sobir1-like* plants. Samples were collected at different timepoints, as indicated, due to varying severity of CD. *Agrobacterium* cultures infiltrated at OD_600_ values of 0.4, 0.8, 0.8 and 0.8 for eGFP-T, INF1-T, INF1 and REL-GFP, respectively. α-Myc-HRP was used to detect PL constructs in input samples, α-GFP-HRP was used to detect REL-GFP and eGFP-T in input and IP samples, and streptavidin-HRP was used to detect biotinylated REL-GFP in IP samples. Arrows indicate biotinylated REL-GFP, red asterisks indicate self-biotinylated INF1-T. Ponceau S staining was used to confirm protein enrichment following GFP-trap bead purification and to verify equal loading of input samples.

In the GFP-IP samples from WT, REL-GFP was detected in all treatments in which it was co-expressed (Figure 5A). As expected, eGFP-T was enriched following immunoprecipitation. Most importantly, streptavidin–HRP detected biotinylated REL only when REL-GFP was co-expressed with INF1-T (Figure 5A). No biotinylated REL was detected in the eGFP-T or INF1 control treatments. Similar results were obtained in both *bak1* and *sobir1/sobir1-like* plants, though longer sampling intervals resulted in more obvious detection of biotinylated REL than in WT plants (Figure 5B, C). Together, these results demonstrate that INF1-T is active when directed to the apoplastic space and specifically biotinylates its interacting receptor REL.

## Discussion

PPI studies are most informative when performed *in vivo*, as this preserves the native context of host–pathogen interactions. Indeed, comparative studies have highlighted significant discrepancies between *in vitro* and *in planta* protein datasets. For instance, Junge and colleagues conducted a study on the systemic secretome of rice leaves both *in planta* and in seed callus suspension-cultured cells (SCCs; *in vitro*). Their findings revealed that only six proteins overlapped between the *in vitro* and *in planta* analyses out of a total of 222 identified proteins (Jung et al. 2008). Moreover, the pathogen *Mycosphaerella fijiensis* was shown to secrete pathogenicity-related proteins in a context-dependent manner, with distinct profiles observed *in vitro* compared with *in planta* infection (Escobar-Tovar et al. 2015). These findings underscore the limitations of *in vitro* systems and highlight the need for *in planta* approaches to achieve an accurate understanding of apoplastic defence responses.

The leaf apoplast differs substantially from the intracellular environment and presents several challenges for PL. It is a highly dynamic compartment involved in water movement and gas exchange, which may influence the stability of apoplastic protein interactions (Gentzel et al. 2019). Its acidic pH (approximately 4.5–5.5; Barbez et al. 2017) also differs markedly from the cytosol (approximately 7; Pittman 2012; Sze and Chanroj 2018) and could affect TurboID activity. Although TurboID remains functional under acidic conditions in the mammalian endoplasmic reticulum lumen (May et al. 2020), performance in the plant apoplast had not been validated. Furthermore, the abundance of apoplastic hydrolases, particularly defensive proteases, may compromise fusion protein stability, as reported previously for apoplastic TurboID constructs (Teplova et al. 2021). These features highlight the need to validate and optimise TurboID activity under apoplastic conditions before applying PL to investigate apoplastic PPIs.

The functionality of INF1-T was assessed based on its ability to induce CD in *N. benthamiana* WT leaves. INF1-T retained the ability to trigger CD, comparable to untagged INF1, indicating that the addition of the TurboID tag did not interfere with INF1 recognition or its ability to activate the corresponding immune response (Figure 1C). In contrast, neither XEG1-T nor untagged XEG1 induced visible CD in *N. benthamiana*. This phenotype may be due to a lack of recognition or perception of XEG1 constructs in the *N. benthamiana* apoplast. A similar functional validation strategy was employed by Zhang et al. (2019), who assessed the functionality of N-TIR-TurboID based on its ability to induce CD. This approach is consistent with the strategy used here to determine whether the addition of TurboID affected the biological activity of INF1.

eGFP-T was detectable when expressed from both vectors, although its accumulation was noticeably higher when expressed from pEAQ3. In contrast, XEG1-T and INF1-T were only detectable in pEAQ3 samples. These results suggest that the pEAQ3 vector generally achieves higher expression levels than pEG101, a pattern previously reported by Kettles et al. (2017) and in studies of the *Z. tritici* Zt6 effector (Kettles et al. 2018). This is likely attributable to the presence of both P19 and the CPMV-HT within the backbone of pEAQ3 (Sainsbury et al. 2009). Moreover, TurboID itself appears to confer a degree of protection against degradation and to improve protein stability (Bell et al. 2013), which could explain why INF1 was only detectable when fused to TurboID.

To allow comparison between treatments, it is essential to achieve comparable accumulation of eGFP-T (control) and INF1-T. GFP has been reported to account for up to 30% of soluble protein following Agroexpression in *N. benthamiana* (Mardanova et al. 2017). Therefore, the relative abundance of eGFP-T and INF1-T was optimised. Increasing the *Agrobacterium* infiltration density increased accumulation of both eGFP-T and INF1-T (Figure 2), consistent with Shamloul et al. (2014). However, INF1-T abundance decreased at 72 hpi, likely due to CD, whereas eGFP-T continued to accumulate. Comparable abundance was achieved at 48 hpi using an OD_600_ of 0.4 for eGFP-T and 0.8 for INF1-T. In future studies, specific conditions will need to be optimised for each target protein prior to performing labelling reactions.

Since INF1-triggered CD interferes with PL, we explored several strategies to attenuate this response. Recognition of some effectors is light-dependent, for example, Su et al. (2018) showed that MPK3 and MPK6 are critical for ETI-associated CD under light conditions, whereas darkness can weaken or abolish this response. We therefore tested whether INF1 recognition was light-dependent by maintaining plants in complete darkness for three days following agroexpression. However, CD developed at a comparable rate under both light and dark conditions (Figure S2), indicating that INF1 recognition is not light-dependent. Calcium signalling is also important for CD induction (Orrenius et al. 2015; Ren et al. 2021). Treatment with 1 mM LaCl_3_ did not reduce INF1-T-induced CD in WT plants compared with the untreated control (Figure S3A). This contrasts with previous reports showing that LaCl_3_ inhibited RPM1-and ZAR1-mediated CD in *A. thaliana* and *N. benthamiana*, respectively (El Kasmi et al. 2017; Hu et al. 2020). This may be explained by mechanistic differences in the induction of CD triggered by cell surface and intracellular immune sensors.

Accurate subcellular localisation of both bait and control proteins is essential for PL because of the promiscuous labelling activity of TurboID. To minimise non-specific biotinylation, both proteins should occupy the same subcellular compartment (Mair et al. 2019). Accordingly, apoplastic localisation of eGFP-T was confirmed by confocal microscopy and compared with cytosolic GFP (Figure S6A). To verify bait localisation, AWF was collected from agroinfiltrated *sobir1*/*sobir1*-like plants. AWF is widely used to investigate the localisation of apoplastic proteins (Sueldo et al. 2024; Hu et al. 2021). Although eGFP-T was readily detected in AWF, INF1-T remained undetectable even after concentration (Figure S6B). However, infiltration of INF1-T-containing AWF into *N. benthamiana* WT plants induced robust CD (Figure S6C), indicating that biologically active INF1-T had accumulated in the apoplast below the western blot detection limit. Alternatively, the Myc tag may have been proteolytically cleaved following secretion, thus preventing detection with anti-Myc antibody whilst retaining INF1-T activity. By contrast, Matsui et al. (2024) detected the apoplastic subtilase SBT5.2 in agroinfiltrated *N. benthamiana* leaves by blotting, suggesting that the inability to detect INF1-T reflects protein-specific properties rather than technical limitations of the AWF approach.

Following confirmation of construct expression and apoplastic localisation, we investigated the feasibility of TurboID-mediated biotinylation in the apoplast. Supplying biotin and ATP exogenously may provide greater control over the labelling reaction by allowing these substrates to be introduced at defined concentrations or timepoints. Previous studies have generally re-infiltrated agroinfiltrated leaves with biotin alone (Li et al. 2023; Matsui et al. 2024; Zhang et al. 2019). However, we supplied biotin, ATP and Mg²⁺ exogenously, which may provide greater control over TurboID-mediated labelling in the apoplast by regulating substrate availability, including the duration and extent of labelling, and may reduce background labelling compared with intracellular expression (Zhang et al. 2019; Teplova et al. 2021). This was particularly relevant for ATP and Mg²⁺, as ATP is required by TurboID to generate the reactive biotin intermediate, yet its concentration in the apoplast is approximately 100-fold lower than inside the cell (Song et al. 2006; Rieder and Neuhaus 2011). Mg²⁺ is also required for ATP binding and biotin-AMP formation (Branon et al. 2018) and is present at relatively low concentrations in the apoplast (Gabriel and Kesselmeier, 1999). The inclusion of both was therefore consistent with previous apoplast-targeted TurboID studies (Teplova et al. 2021; Mismar et al. 2024).

Western blotting using streptavidin-HRP confirmed that TurboID-mediated biotinylation occurred successfully in the apoplast (Figure 3). Furthermore, the biotinylation profile generated by INF1-T differed from that of the eGFP-T control, consistent with previous TurboID studies (Li et al. 2023; Matsui et al. 2024), while self-biotinylation of both constructs was also observed, as expected (Zhang et al., 2019).

Co-IP further demonstrated specific biotinylation of REL-GFP by INF1-T in WT, *bak1* and *sobir1 N. benthamiana* backgrounds (Figure 5). REL-GFP biotinylation occurred exclusively in the presence of INF1-T and was not detected with the eGFP-T control, despite the higher accumulation of eGFP-T and overexpression of REL-GFP. These findings demonstrate that TurboID-mediated PL can achieve specific protein biotinylation within the apoplast. Chen et al. (2023) previously reported the interaction between INF1 and REL in WT *N. benthamiana* using Co-IP. In the present study, this interaction was successfully detected in both *bak1* and *sobir1* mutant backgrounds, demonstrating that disruption of these co-receptors does not prevent the association between INF1 and REL from being captured using a TurboID-based PL approach. To our knowledge, this is the first report demonstrating successful PL of the INF1–REL interaction in these mutant backgrounds.

Application of this method as a screening tool to identify unknown elicitor-receptor interactions will be desirable in the future. Demonstrating feasibility in a CD-inducing interaction provides stringent validation, as CD interferes with downstream biochemical analyses. Although future studies may be better suited to studying interactions that do not trigger CD, our results demonstrate that this limitation can be circumvented using *bak1* or *sobir1* mutant backgrounds. Thus, the approach remains applicable to CD-inducing apoplastic effectors, while potentially being more straightforward in systems where CD is not triggered. To our knowledge, this is the first study to establish and validate a TurboID-based PL system for investigating extracellular protein interactions in the plant apoplast. This finding is consistent with the recent study by Mismar et al. (2024), who demonstrated the feasibility of a similar TurboID-based strategy in the extracellular environment of mammalian cells, highlighting the broader applicability of extracellular PL across biological systems.

## Materials and methods

### Plant growth conditions

*N. benthamiana* seeds were sown in a single 11 cm pot containing water-saturated Jiffy peat-free compost supplemented with Osmocote fertiliser (2.5 g per pot). The pot was kept fully submerged in water until seed germination. Germination and early growth were carried out in a controlled growth room under long-day conditions (16 h light/ 8 h dark photoperiod) at 20 °C and relative humidity above 80% for two weeks. Following this period, individual seedlings were transplanted into separate 11 cm pots filled with fresh Jiffy compost and supplemented again with 2.5 g Osmocote fertiliser per pot.

### Bacterial growth conditions

All *E. coli* and *Agrobacterium tumefaciens* strains (GV3101) were cultured in Luria– Bertani (LB) Lennox medium, used as broth for liquid cultures and solidified with agar for plates. The 70 ml liquid cultures in 200 ml baffled conical flask were incubated in in a shaking incubator at 200 rpm. *E. coli* strains were grown overnight at 37 °C, whereas *A. tumefaciens* strains were incubated at 28 °C for 36–48 h.

### Cloning of PL constructs

The native SPs of XEG1 and INF1 were predicted using SignalP (Almagro Armenteros et al. 2019) and replaced with the NbPR1 SP. PL constructs eGFP-T, XEG1-T, XEG1, INF1-T, and INF1, as listed in Table S1, were synthesised by Twist Bioscience (San Francisco, USA) for expression in the binary vector pEAQ3 (Sainsbury et al. 2009) and pEG101 (Earley et al. 2006) subsequently cloned using Gateway recombination.

The constructs XEG1 and INF1 were synthesised as gene fragments and amplified by PCR to introduce attB sites using Phusion High-Fidelity DNA Polymerase and attB-flanked primers (Table S2). The PCR programme consisted of 95 °C for 30 s, followed by 30 cycles of 95 °C for 30 s, 74 °C for 30 s, and 72 °C for 1 min, with a final extension at 72 °C for 5 min. Products were gel-purified using the QIAquick Gel Extraction Kit. BP recombination reactions were performed to transfer PCR products into pDONR207. The reaction was incubated overnight at room temperature (RT) and terminated with Proteinase K. Recombinant plasmids were transformed into *E. coli* NEB 10-beta by electroporation, selected on LB–gentamicin plates, and verified by colony PCR (attL primers, Table S2). Colony PCR was performed using the same cycling conditions described above, except that the annealing temperature was 66 °C and 30 cycles were used. For constructs synthesised in pTwist Entry vector (eGFP-T, XEG1-T, and INF1-T), inserts were PCR-amplified using M13 primers (Table S2) with Phusion polymerase and purified as above. PCR conditions were identical to those described above, except that the annealing temperature was 58°C. All constructs were transferred into pEAQ3 and pEG101 using LR recombination. The LR reaction was incubated overnight at RT, terminated with Proteinase K, and transformed into *E. coli* NEB 10-beta. Transformants were selected on LB– kanamycin plates and confirmed by colony PCR (pEAQ3 and pEG101 primers; Table S2). Colony PCR was performed using the cycling conditions described above, with annealing temperatures of 72 °C for pEAQ3 primers and 66 °C for pEG101 primers, and 30 amplification cycles.

Plasmids were extracted using the QIAprep Spin Miniprep Kit and verified by full sequencing (PlasmidsNG, Birmingham, UK). Confirmed constructs were transformed into *A. tumefaciens* GV3101 by electroporation, selected on LB plates containing gentamicin and kanamycin, and incubated at 28 °C for 48 h. Glycerol stocks were stored at −80 °C.

### *A. tumefaciens* transient expression

*Agrobacterium*-mediated transient expression was performed as previously described (Kapila et al. 1997). *A. tumefaciens* cultures were grown in 70 ml LB supplemented with 25 µg/ml gentamicin and 50 µg/ml kanamycin in 250 ml baffled Erlenmeyer flasks. Cultures were pelleted by centrifugation at 8,500 rpm for 3 min at RT. The supernatant was discarded and the cells were washed twice by resuspension in agroinfiltration buffer (10 mM MgCl₂, 10 mM MES, pH 5.6) followed by centrifugation under the same conditions. To determine density, 100 µl of each final cell suspension was diluted in 900 µl buffer, and the OD_600_ was measured using a spectrophotometer. Cell suspensions were then diluted to the required OD_600_ using buffer. Acetosyringone was added to a final concentration of 150 µM, and the suspensions were mixed gently by inversion before incubation in the dark for 2–3 h prior to infiltration. Leaves of 4–6-week-old *N. benthamiana* were infiltrated with the *Agrobacterium* suspensions using 1 ml needleless syringes.

### Assessment of CD phenotypes

To assess CD, five-week-old *N. benthamiana* plants were agroinfiltrated with the appropriate *Agrobacterium* strains at OD_600_ = 0.7. CD phenotypes were assessed at 4 dpi by visual inspection and photography. CD was further quantified by measuring red fluorescence (317 nm; arbitrary units, AU) from 20 leaves per treatment, as previously described by Xi et al. (2021) using a Typhoon scanner.

### Total protein extraction

Following *Agrobacterium*-mediated transient expression, agroinfiltrated leaves were harvested at the required time point post-infiltration. The central vein of each leaf was removed, and the fresh weight of two infiltrated leaves per plant was recorded. Leaf tissue was wrapped in aluminium foil, snap-frozen in liquid nitrogen, and stored at −80 °C until further processing. For protein extraction, frozen leaf tissue was ground to a fine powder using a sterile mortar and pestle pre-cooled with liquid nitrogen. The resulting powder was transferred to 15 ml Falcon tubes and maintained in liquid nitrogen throughout sample handling. The base extraction buffer (GTEN) was prepared in advance as previously described by Sarris et al. (2015) and consisted of 150 mM Tris–HCl (pH 7.5), 150 mM NaCl, 1 mM ethylenediaminetetraacetic acid (EDTA), and 10% (v/v) glycerol, with distilled water (dH_2_O) added to the final volume. The buffer was stored at 4 °C until use. On the day of protein extraction, a fresh working extraction buffer was prepared by supplementing the GTEN buffer with 1% (v/v) cOmplete™ EDTA-free protease inhibitor cocktail (Sigma–Aldrich), 1% (v/v) IGEPAL, 2% (w/v) polyvinylpolypyrrolidone (PVPP), 0.5% (w/v) sodium deoxycholate, 10 mM dithiothreitol (DTT), and 0.1% (w/v) sodium dodecyl sulfate (SDS). Three volumes of the working extraction buffer, relative to the fresh weight of the leaf tissue, were added to each sample. Samples were vortexed thoroughly and incubated on a tube rotator at 4 °C for 20 min. The extracts were then centrifuged at 4,000 rpm for 25 min at 4 °C to pellet insoluble debris. The resulting supernatant, containing the total soluble protein fraction, was carefully transferred to fresh tubes and used for subsequent analyses.

### AWF extraction

AWF extraction was performed using a harvest buffer prepared according to Kingsbury and McDonald (2014), with minor modifications. The buffer consisted of 20 mM sodium acetate (pH 5.5) and 100 mM NaCl. At the required time point, the harvest buffer was pre-chilled to 4 °C and infiltrated into *N. benthamiana* leaves using a 1 ml needleless syringe. Following infiltration, leaves were excised and gently rolled around a 1 ml blue pipette tip before being placed into a 20 ml syringe without the plunger. The syringe was then positioned inside a 50 ml Falcon tube and kept on ice to minimise protein degradation. Samples were centrifuged at 4,000 rpm for 10 min at 4 °C, and the AWF was collected from the bottom of each 50 ml Falcon tube. Aliquots of the AWF were taken for SDS–PAGE and protein accumulation analysis, while a portion was reinfiltrated into *N. benthamiana* leaves for further assays. For concentration of the AWF, Amicon Ultra centrifugal filter units with a 3 kDa molecular weight cut-off (MWCO) were used, followed by centrifugation at 4 °C for 20 min.

### SDS-PAGE and Western blots

Protein samples for SDS–PAGE were prepared by mixing protein extracts with 3x SDS loading buffer containing 10% (v/v) DTT at a ratio of 2:1 (sample: buffer). Samples were vortexed and denatured at 95 °C for 5 min at 300 rpm using an Eppendorf ThermoMixer. Polyacrylamide gels were prepared according to the Cold Spring Harbor Laboratory protocol (CSH Press, 2015). Resolving gels of 10%, 12%, and 15% acrylamide were cast. A 4% stacking gel was then prepared according to the CSH protocol (CSH Press, 2015), poured on top of the resolving gel, and a comb was inserted to form wells. Where indicated, 4–20% Mini-PROTEAN TGX pre-cast gels (12-well; Bio-Rad) were used for analysis of biotinylated proteins. Following denaturation, samples were loaded into the wells alongside 6 µl of pre-stained protein ladder (PageRuler Plus, Thermo Scientific; 10–170 kDa). Electrophoresis was performed using a Bio-Rad Mini-PROTEAN Tetra system. Gels were initially run at 80 V for 15–20 min until samples entered the resolving gel, after which the voltage was increased to 100–120 V and electrophoresis continued at RT until completion. Following electrophoresis, proteins were transferred onto Amersham Protran 0.45 µm nitrocellulose membranes using a wet transfer system (Bio-Rad Mini-PROTEAN Tetra) at 30 V overnight at 4 °C in transfer buffer (25 mM Tris, 192 mM glycine, 10% methanol; pH 8.3). Following transfer, membranes were briefly rinsed in dH_2_O and blocked for 1 h at RT with gentle agitation. Membranes used for Myc-or GFP-tag detection were blocked in 5% (w/v) skimmed milk prepared in Tris-buffered saline containing 0.1% Tween-20 (TBST), whereas membranes used for streptavidin–HRP detection were blocked in 2% (w/v) bovine serum albumin (BSA) in TBST. Tris-buffered saline (TBS) consisted of 20 mM Tris and 150 mM NaCl (pH 7.6). Membranes were then incubated overnight at 4 °C with the appropriate primary antibody or probe: α-Myc–HRP or α-GFP–HRP (1:500 dilution in 5% milk/TBST), or streptavidin–HRP (1:10,000 dilution in 2% BSA/TBST). The following day, membranes were washed three times with TBST for 5 min each at RT. Protein detection was performed using SuperSignal West Pico PLUS Chemiluminescent Substrate (Thermo Scientific), and signals were visualised using an Amersham Imager 680 (GE Healthcare).

### *In planta* biotinylation assay

Due to limited availability of essential cofactors for the biotinylation reaction in the *N. benthamiana* apoplast (ATP and Mg^2+^) an *in planta* biotinylation approach was used as previously described by Teplova et al. (2021), with minor modifications. The biotinylation solution consisted of 200 µM biotin, 500 µM ATP, and 1.2 mM magnesium acetate, and was prepared freshly prior to use. At the appropriate time points, agroinfiltrated leaves were infiltrated with the biotinylation solution using a 1 ml needleless syringe. Leaves were then incubated at RT for 1–3 h to allow the biotinylation reaction to proceed before harvesting.

### Streptavidin bead pull-down assay

This assay was performed according to Szymansky (2025), with minor modifications. Following the biotinylation assay and total protein extraction as described above, residual free biotin was removed from the protein supernatant using Zeba Spin Desalting Columns (7K MWCO, 10 ml; Thermo Scientific). The protein extracts were first filtered through Miracloth (Merck Millipore) and then desalted by centrifugation at 1,000 x *g* for 2 min at 4 °C. The desalted protein eluates were collected into pre-chilled 15 ml Falcon tubes. High-capacity streptavidin agarose beads (Pierce, Thermo Scientific) were added to the desalted protein extracts (400 µl of a 50% bead slurry, equivalent to 200 µl bead volume) and incubated for 2 h at 4 °C on a rotating mixer to allow binding of biotinylated proteins. Following incubation, the beads were pelleted by centrifugation at 3,000 rpm for 2 min at 4 °C and transferred to 1.5 ml microcentrifuge tubes. The beads were washed twice with GTEN buffer, followed by one wash with 7 M urea prepared in 50 mM ammonium bicarbonate (pH 8.0), two washes with 4 M urea prepared in 50 mM ammonium bicarbonate (pH 8.0), and three washes with ice-cold 100 mM ammonium bicarbonate buffer to remove non-specifically bound proteins. After the final wash, the beads were resuspended in 1 ml of ice-cold 50 mM ammonium bicarbonate buffer. Aliquots were collected for SDS– PAGE analysis, and the remaining bead suspensions were stored at −80 °C for downstream analyses. Streptavidin bead eluates were separated on 4–20% Mini-PROTEAN TGX pre-cast gradient gels (Bio-Rad), and biotinylated proteins were detected by western blotting using streptavidin–HRP.

### Co-IP assay

This experiment was performed to confirm the biotinylation of the cell surface RLP NbREL (NbL18g11150.1) following its interaction with INF1-T in the presence of biotin. NbREL was fused to GFP in the binary vector SOL2095 (REL-GFP; Bi et al., 2016) and co-expressed with INF1-T, eGFP-T, or INF1. *Agrobacterium* strains were infiltrated at final OD_600_ values of 0.8 for REL-GFP, INF1-T, and INF1, and 0.4 for eGFP-T. Agroinfiltration was performed in WT, *sobir1* (Huang et al. 2021), and *bak1* (Sun et al. 2022) *N. benthamiana* plants. Biotin was infiltrated approximately 1 h before sample collection. Leaf tissue was harvested at 36 hpi for WT, 60 hpi for *sobir1*, and 46 hpi for *bak1*. For each treatment, 2 g of infiltrated leaf tissue was collected. Protein extraction was performed as described above. Cleared protein extracts were filtered through Miracloth (Merck Millipore), and input samples were collected prior to immunoprecipitation. GFP-tagged proteins were immunoprecipitated using GFP-Trap agarose beads (ChromoTek, Proteintech; 19 µl bead slurry per sample). The beads were pre-equilibrated with GTEN buffer and incubated with the protein extracts for 3 h at 4 °C on a rotating mixer. Following incubation, the beads were washed three times with GTEN buffer supplemented with 1 mM DTT, 1x protease inhibitor cocktail, and 0.5% IGEPAL. After the final wash, the beads were resuspended in 30 µl of washing buffer, mixed with 15 µl of 3x SDS loading buffer, and the immunoprecipitated proteins were eluted for SDS–PAGE and western blot.

## Statistical analysis

All statistical analyses were performed using R version 4.3.1 (R Core Team, 2023). Data are presented as the mean ± standard deviation (SD). Pairwise comparisons were performed using Student’s *t*-test. Statistical analyses were conducted using the car, multcomp, stats, and dplyr packages in R.

## Acknowledgements

We would like to thank Prof. Jonathan Jones for hosting AMQ for a short visit to his group at The Sainsbury Laboratory (TSL) and for providing training in the TurboID-based proximity labelling technique. We also thank Prof. Cyril Zipfel (TSL) and Prof. Yuanchao Wang (Nanjing Agricultural University) for providing the *bak1* seeds, and Dr Matthieu Joosten (Wageningen University) for providing the *sobir1 & sobir1-like* seeds and the REL-GFP construct. We would also like to thank Dr Rory Osborne (University of Birmingham) and Dr Hee-Kyung Ahn (University of Edinburgh) for their valuable assistance with protein and Co-IP work. AMQ was supported by a PhD scholarship at the University of Birmingham, funded by the Newton-Mosharafa Fund and the Egyptian Ministry of Higher Education (Mission Sector). ChatGPT was used to assist with language editing of the manuscript.

## Author Contributions

AMQ designed and performed all experiments, analysed the data and wrote the draft of the manuscript. CMS developed and shared a draft protocol for TurboID labelling in *N. benthamiana*. GJK conceived the project, designed the experiments and edited the manuscript.

