## Supplementary Data for "A TurboID-based proximity labelling method for detecting extracellular protein interactions in plants"

| **Gene name** | **Sequence** |
| --- | --- |
| TurboID | ATGAAAGACAATACTGTGCCTCTGAAGCTGATCGCTCTCCTGGCTAATGGCGAGTTCCATAGTGGCGAACAGCTGGGAGAAACCCTGGGCATGTCCAGGGCCGCTATCAACAAGCACATTCAGACTCTGCGCGACTGGGGCGTGGACGTGTTCACCGTGCCCGGAAAGGGCTACTCTCTGCCCGAGCCTATCCCGCTGCTGAACGCTAAACAGATTCTGGGACAGCTGGACGGCGGGAGCGTGGCAGTCCTGCCTGTGGTCGACTCCACCAATCAGTACCTGCTGGATCGAATCGGCGAGCTGAAGAGTGGGGATGCTTGCATTGCAGAATATCAGCAGGCAGGGAGAGGAAGCAGAGGGAGGAAATGGTTCTCTCCTTTTGGAGCTAACCTGTACCTGAGTATGTTTTGGCGCCTGAAGCGGGGACCAGCAGCAATCGGCCTGGGCCCGGTCATCGGAATTGTCATGGCAGAAGCGCTGCGAAAGCTGGGAGCAGACAAGGTGCGAGTCAAATGGCCCAATGACCTGTATCTGCAGGATAGAAAGCTGGCAGGCATCCTGGTGGAGCTGGCCGGAATAACAGGCGATGCTGCACAGATCGTCATTGGCGCCGGGATTAACGTGGCTATGAGGCGCGTGGAGGAAAGCGTGGTCAATCAGGGCTGGATCACACTGCAGGAAGCAGGGATTAACCTGGACAGGAATACTCTGGCCGCTACGCTGATCCGAGAGCTGCGGGCAGCCCTGGAACTGTTCGAGCAGGAAGGCCTGGCTCCATATCTGCCACGGTGGGAGAAGCTGGATAACTTCATCAATAGACCCGTGAAGCTGATCATTGGGGACAAAGAGATTTTCGGGATTAGCCGGGGGATTGATAAACAGGGAGCCCTGCTGCTGGAACAGGACGGAGTTATCAAACCCTGGATGGGCGGAGAAATCAGTCTGCGGTCTGCCGAAAAG |
| eGFP | ATGGTGAGCAAGGGCGAGGAGCTGTTCACCGGGGTGGTGCCCATCCTGGTCGAGCTGGACGGCGACGTAAACGGCCACAAGTTCAGCGTGTCCGGCGAGGGCGAGGGCGATGCCACCTACGGCAAGCTGACCCTGAAGTTCATCTGCACCACCGGCAAGCTGCCCGTGCCCTGGCCCACCCTCGTGACCACCCTGACCTACGGCGTGCAGTGCTTCAGCCGCTACCCCGACCACATGAAGCAGCACGACTTCTTCAAGTCCGCCATGCCCGAAGGCTACGTCCAGGAGCGCACCATCTTCTTCAAGGACGACGGCAACTACAAGACCCGCGCCGAGGTGAAGTTCGAGGGCGACACCCTGGTGAACCGCATCGAGCTGAAGGGCATCGACTTCAAGGAGGACGGCAACATCCTGGGGCACAAGCTGGAGTACAACTACAACAGCCACAACGTCTATATCATGGCCGACAAGCAGAAGAACGGCATCAAGGTGAACTTCAAGATCCGCCACAACATCGAGGACGGCAGCGTGCAGCTCGCCGACCACTACCAGCAGAACACCCCCATCGGCGACGGCCCCGTGCTGCTGCCCGACAACCACTACCTGAGCACCCAGTCCGCCCTGAGCAAAGACCCCAACGAGAAGCGCGATCACATGGTCCTGCTGGAGTTCGTGACCGCCGCCGGGATCACTCTCGGCATGGACGAGCTGTACAAG |
| INF1 (without native SP) | ACCACGTGCACCACCTCGCAGCAGACCGTAGCGTACGTGGCGCTCGTAAGCATCCTCTCGGACACGTCGTTTAATCAGTGCTCGACGGACTCCGGCTACTCGATGCTGACGGCCACCTCGCTGCCCACGACGGAGCAGTACAAGCTCATGTGCGCGTCGACGGCGTGCAAGACGATGATCAACAAGATCGTGTCGCTCAACGCTCCCGACTGCGAGCTGACGGTGCCAACTAGTGGCCTGGTACTCAACGTGTACTCGTACGCGAACGGGTTCTCGTCTACGTGTGCGTCGCTATGA |
| XEG1 (without native SP) | TACATCGTGTACAACAACCTGTGGAACAAGAACGCTGCTGCTTCTGGTTCTCAGTGCACCGGTGTGGATAAGATCAGCGGTTCTACTATTGCCTGGCACACCTCTTACACTTGGACTGGTGGTGCTGCTACTGAGGTGAAGTCTTACTCTAACGCTGCCCTGGTGTTCAGCAAGAAGCAGATCAAGAACATCAAGAGCATCCCGACCAAGATGAAGTACAGCTACAGCCACTCTTCTGGCACCTTCGTTGCTGATGTGAGCTACGATCTGTTTACCAGCTCTACCGCTTCCGGCTCTAACGAGTACGAGATTATGATTTGGCTGGCTGCTTACGGTGGCGCTGGTCCTATTTCTTCTACCGGTAAGGCTATTGCCACCGTGACCATCGGTAGCAACAGCTTCAAGCTTTACAAGGGCCCTAACGGTAGCACCACCGTTTTCTCATTCGTGGCTACCAAGACCATCACCAACTTCTCTGCTGACCTGCAGAAGTTCCTGAGCTACCTTACTAAGAACCAGGGCCTGCCATCTAGCCAGTACCTTATTACTCTTGAGGCTGGCACTGAGCCTTTCGTGGGTACTAATGCTAAGATGACCGTGTCCAGCTTCAGCGCTGCTGTGAAT |
| N-terminal fusion linker | GGTTCGATGGCTCGGGATCCACCGGTCGCCACC |
| C-terminal fusion linker | GCTAGC |

**Table S1**. DNA sequences of the PL constructs used in this study.

**Table S2**.PCR primers for PL cloning.

| **Primer names** | **Sequences** |
| --- | --- |
| M13-F | GTAAAACGACGGCCAG |
| M13-R | CAGGAAACAGCTATGAC |
| attB1-F | GGGGACAAGTTTGTACAAAAAAGCAGGCTTAATGGGATTTGTTCTTTTTTCTC |
| attB2-R | GGGGACCACTTTGTACAAGAAAGCTGGGTATTACAGATCCTCTTCTGAGA |
| attL1-F | GCAGGCTATGGGATTTGTTCTTT |
| attL2-R | GAAAGCTGGGTTTACAGATCCTCT |
| pEG101-F | TCGCAAGACCCTTCCTCTAT |
| pEG101-R | TACGTCGCCGTCCAGCTCGA |
| pEAQ3-F | AACGTTGTCAGATCGTGCTTCGGCACC |
| pEAQ3-R | CTGAAGGGACGACCTGCTAAACAGGAG |

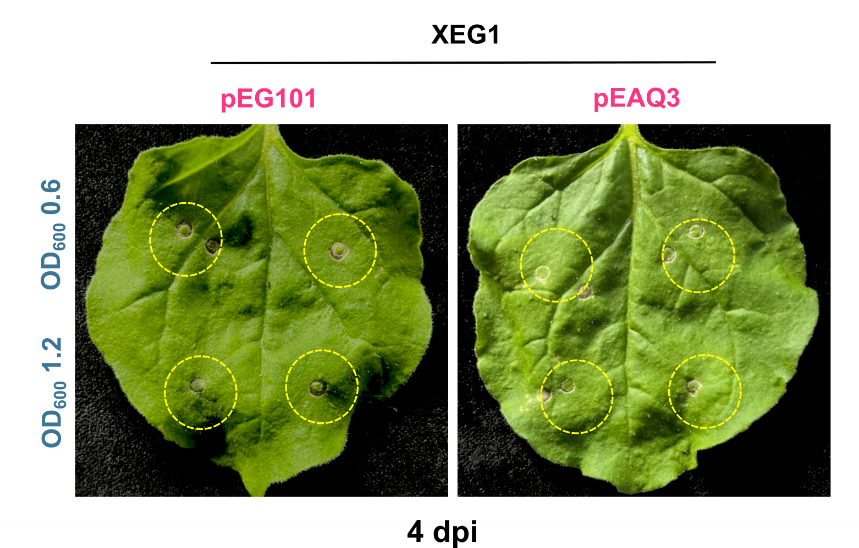

**Figure S1. XEG1 does not induce CD in N. benthamiana WT plants.** Agrobacterium-mediated transient expression of the XEG1 construct was performed at OD_600_ values of 1.2 and 0.6 using the pEAQ3 and pEG101 vectors. The ability of XEG1 expressed from either vector to induce a CD response was assessed at 4 dpi. The experiment was repeated twice with comparable results.

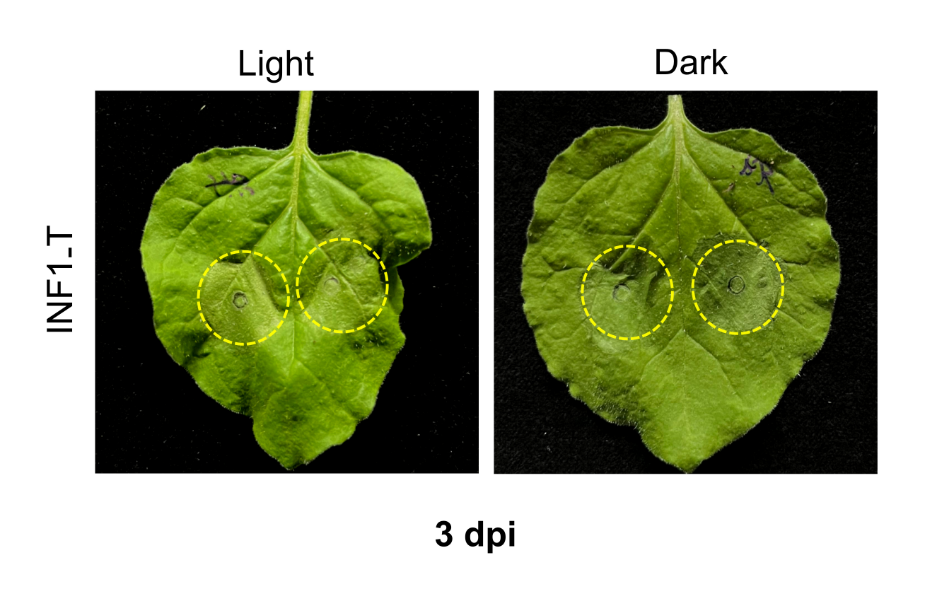

**Figure S2. Dark treatment did not affect the CD response to INF1-T.** To investigate whether the CD response to INF1-T is light dependent, N. benthamiana WT leaves were agorinfiltrated with INF1-T (OD_600_ = 0.8). Agroinfiltrated leaves were either kept in the dark immediately after infiltration or maintained under normal light-dark cycles as a control. The cycles indicate the agroinfiltrated area within the leaf. The CD response was assessed at three dpi. The experiment was repeated twice with comparable outcomes.

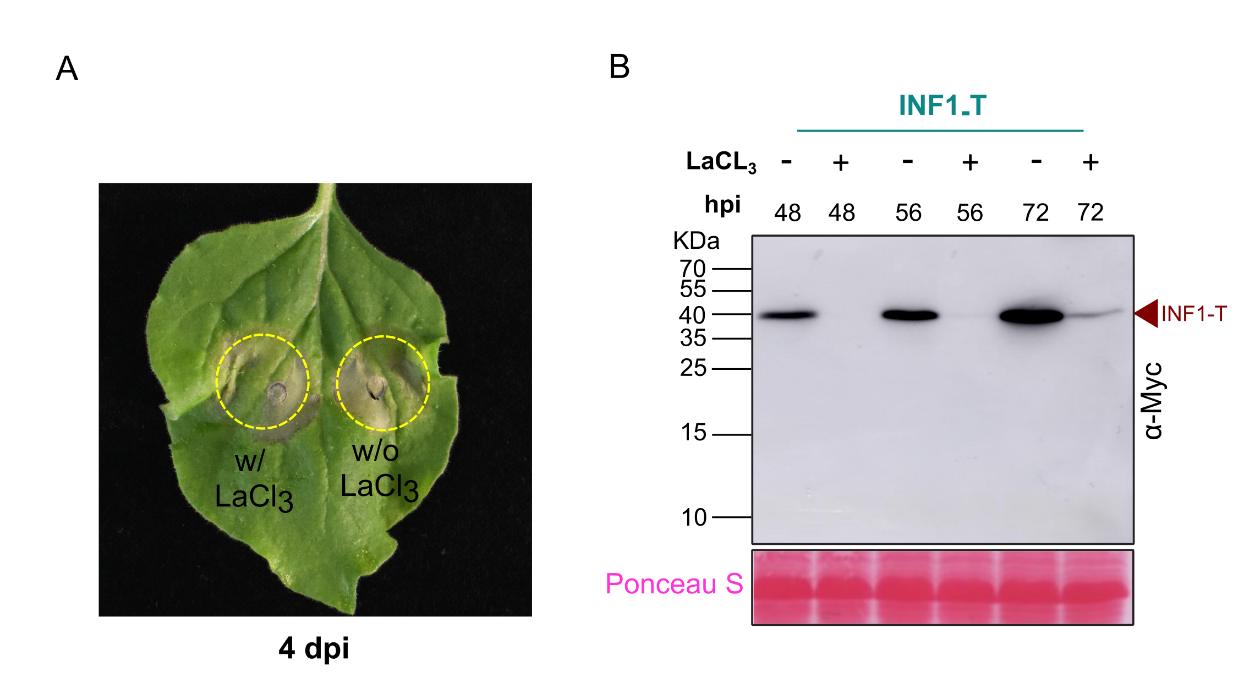

**Figure S3. Calcium channel blocker (LaCl_3_) did not affect the CD phenotype development.** (A) CD phenotype development in N. benthamiana WT leaves following agroinfiltration of INF1-T (OD_600_ = 0.8) with or without 1 mM LaCl3. LaCl_3_ was co-infiltrated with the Agrobacterium strain on the left side of the leaf, whereas INF1-T was infiltrated without LaCl_3_ on the right side. The CD phenotype was assessed at 4 dpi. (B) INF1-T accumulation was analysed by western blotting using α-Myc-HRP in samples with or without co-infiltration of 1 mM LaCl_3_ at 48, 56 and 72 hpi. Equal protein loading was confirmed by Ponceau S staining.

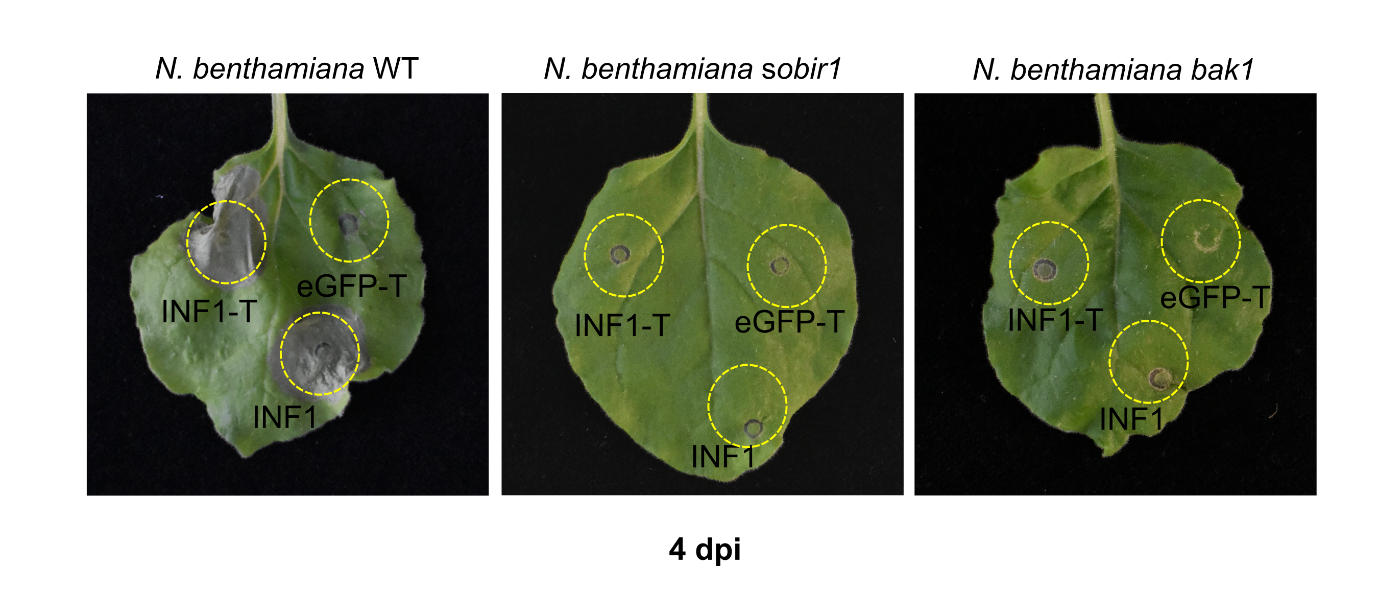

**Figure S4. INF1-induced CD is abolished in sobir1/sobir1-like and bak1 plants.** WT plants were used as a control. The CD phenotype induced by INF1 and INF1-T was assessed at 4 dpi. Agrobacterium cultures were infiltrated at OD_600_ values of 0.4 for eGFP-T and 0.8 for INF1 and INF1-T.

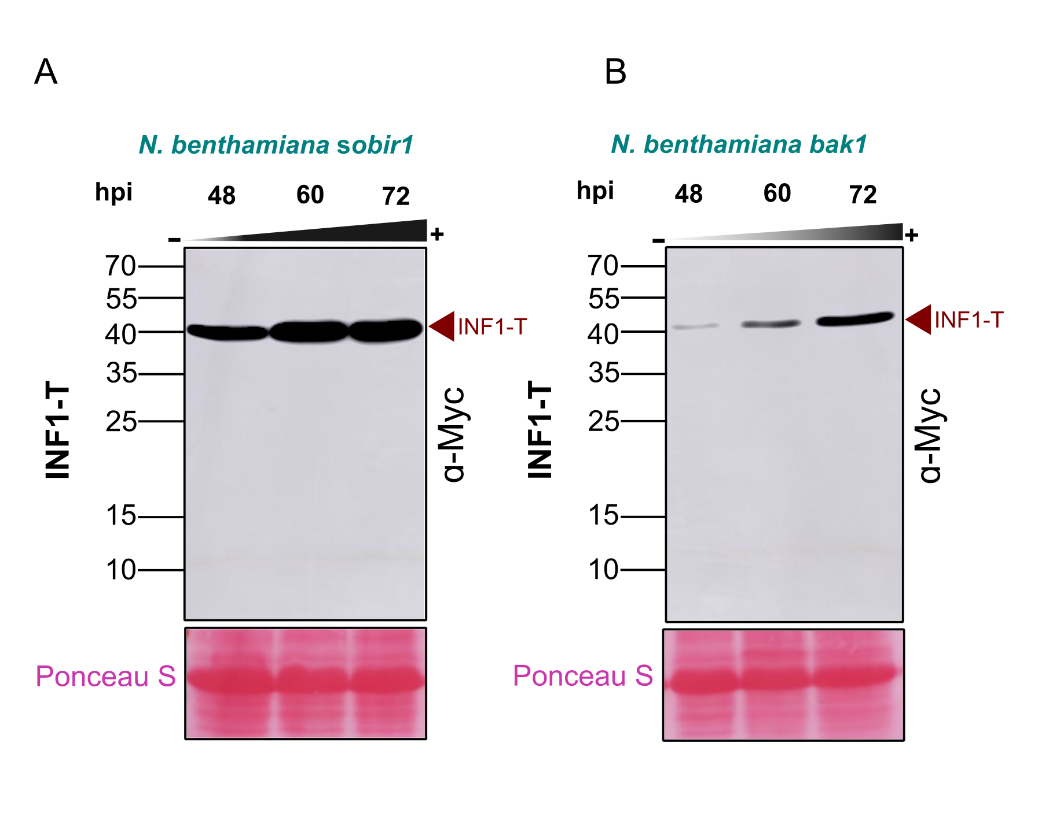

**Figure S5. The abundance of INF1 constructs in sobir1/sobir1-like and bak1 plants.** Protein accumulation of INF1-T across three time points (48, 60 and 72 dpi) in the indicated genotypes. Western blotting with α-Myc HRP was used to assess protein levels, showing that INF1-T gradually accumulated over time in sobir1/sobir1-like and bak1 plants.

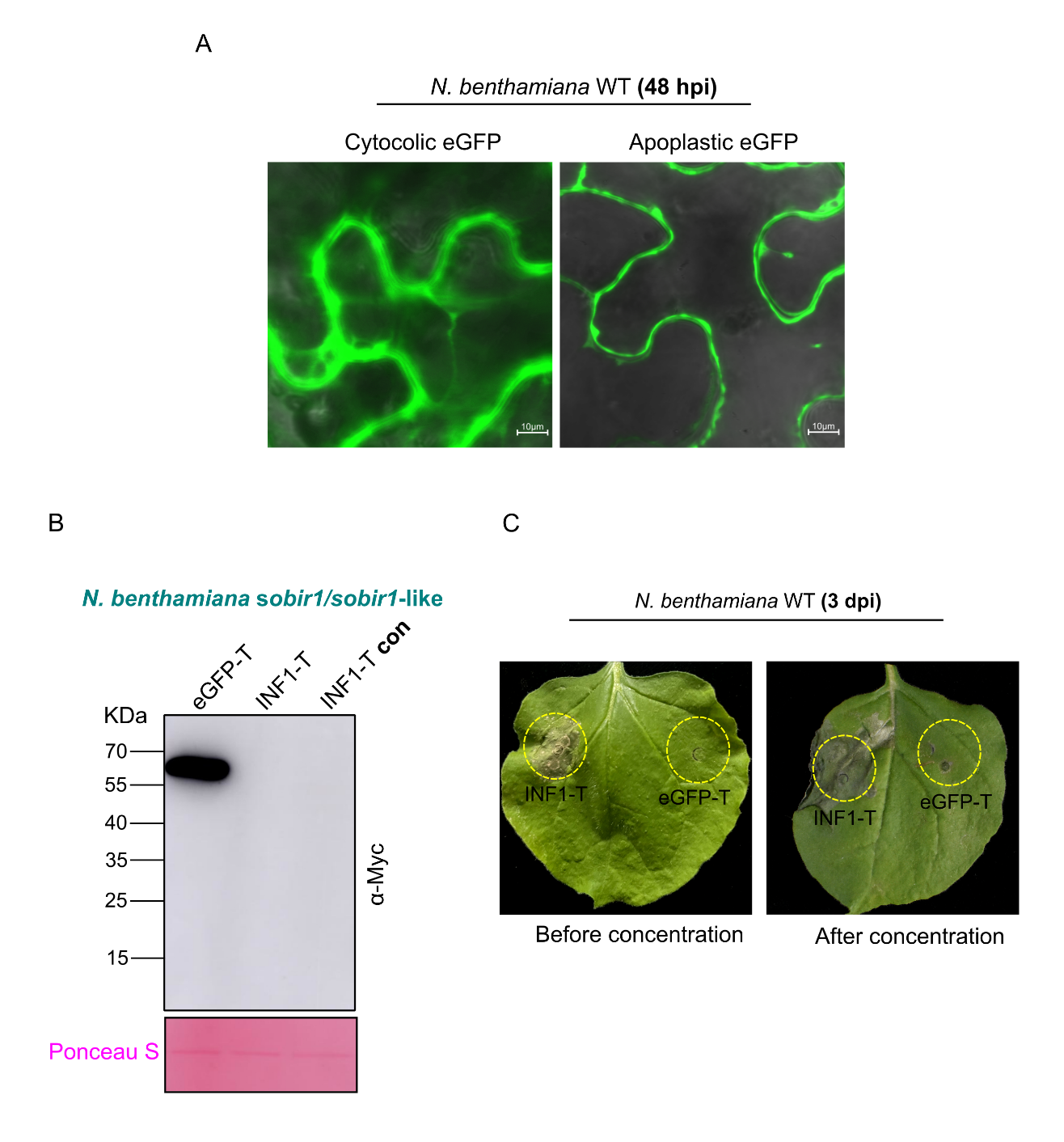

**Figure S6. eGFP-T and INF1-T localise to the N. benthamiana apoplast.** (A) Apoplastic localisation of eGFP-T was indicated by fluorescence restricted to the cell periphery, in contrast to the cytosolic GFP control. Images were captured using Zeiss LSM 880 confocal microscopy at 48 hpi. Agrobacterium cultures expressing GFP-T (+NbPR1 SP) or cytosolic GFP (−NbPR1 SP) from the pEAQ3 vector were infiltrated at an OD_600_ value of 0.4. (B) Detection of eGFP-T and INF1-T in AWF by western blotting using α-Myc-HRP. AWF was collected at 72 hpi from sobir1/sobir1-like leaves and concentrated using centrifugal filter units. Potential contamination with cytosolic proteins was assessed by Ponceau S staining. (C) AWF from each construct, before and after concentration, was re-infiltrated into N. benthamiana WT leaves. The CD phenotype was assessed at 3 dpi.

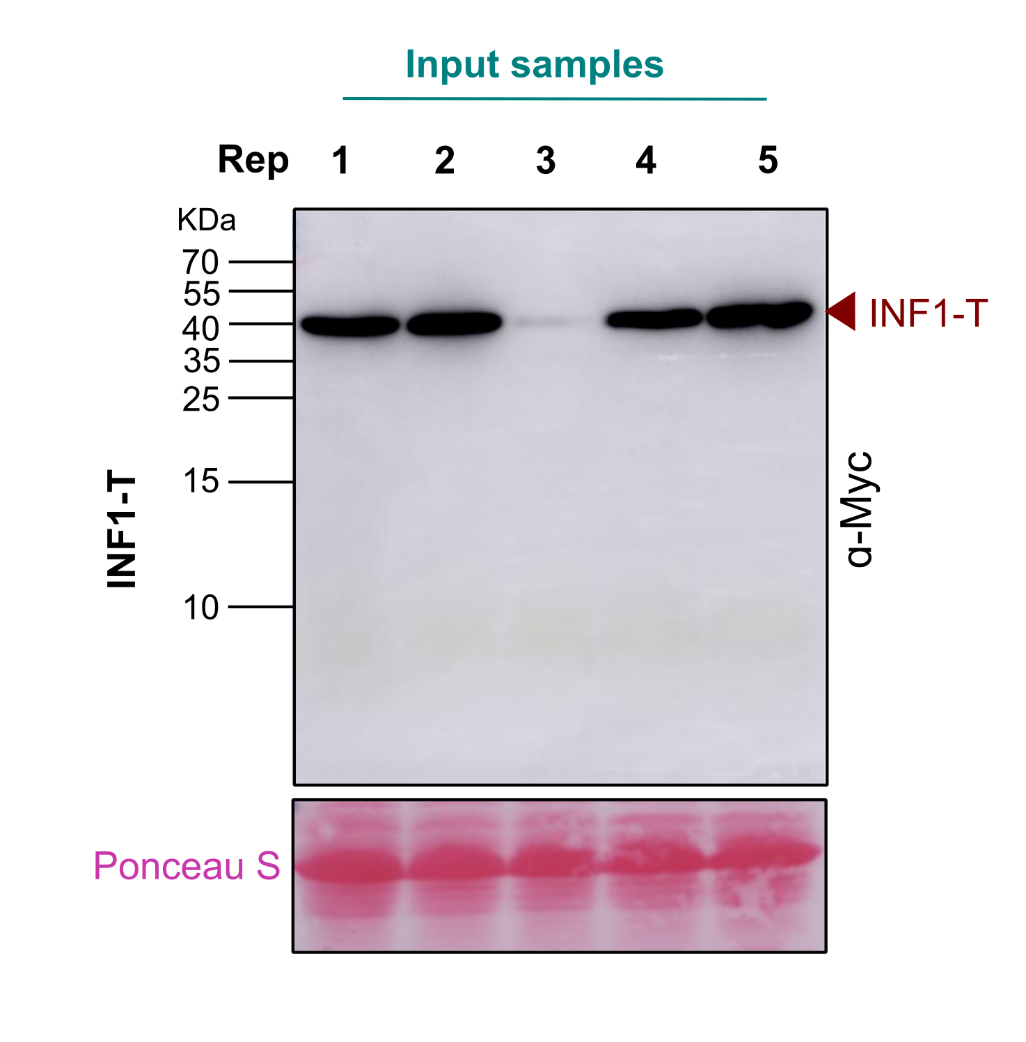

**Figure S7. NF1-T accumulation in input samples following the biotinylation assay.** INF1-T accumulation in N. benthamiana WT plants was analysed by Western blotting using α-Myc-HRP following the biotinylation assay and total protein extraction. The expected INF1-T band was detected in 4 out of 5 biological replicates (indicated by the red arrow). Equal protein loading was confirmed by Ponceau S staining.
